# Population and conservation genomics of the world's rarest hyena species, the brown hyena (*Parahyena brunnea*)

**DOI:** 10.1101/170621

**Authors:** Michael Westbury, Stefanie Hartmann, Axel Barlow, Ingrid Wiesel, Viyanna Leo, Rebecca Welch, Daniel M Parker, Florian Sicks, Arne Ludwig, Love Dalén, Michael Hofreiter

## Abstract

With an estimated population size of less than 10,000 individuals worldwide, the brown hyena (*Parahyaena brunnea*) has been listed as ‘near threatened’ by the IUCN. Despite this rank, studies involving DNA analyses of the brown hyena are limited. Little consideration has been focussed towards population structure within the brown hyena, which could provide valuable insights about its evolutionary history and aid in conservation efforts of the species. Here we report both mitochondrial and nuclear genomes of wild-caught brown hyena individuals from across southern Africa. Mitochondrial DNA shows little to no phylogeographic structure, whereas low-coverage nuclear genomes reveal several potential sub-populations. Moreover, we find that brown hyenas harbour the lowest genetic diversity for a species on both the mitochondrial and nuclear level when compared to a number of mammalian species for which such information is currently available. Our data also reveal that at least on the nuclear DNA level, this low diversity could be the result of a continuous and ongoing decline in effective population size that started about one million years ago and dramatically accelerated towards the end of the Pleistocene. Moreover, our findings also show that the correlation between genetic diversity and the perceived risk of extinction is not particularly strong, since many species with higher genetic diversity than the brown hyena are considered to be at greater risk of extinction. Taken together, our results have important implications for the conservation status and conservation approaches of the brown hyena.

## Introduction

With the ever-decreasing price of next generation sequencing (NGS), population genomics has become a rapidly evolving field within evolutionary biology (Buerkle and Gompert 2013). First appearing in the late 1990s with the analysis of large-scale datasets on human single-nucleotide-polymorphisms (Hartl and Clark 1997), population genomics analyses have now been implemented in the study of a range of species and has greatly aided in understanding their evolution (Ellegren 2014; Liti et al. 2009; Nielsen et al. 2009). A major argument in favour of the analysis of whole genomic data is that the high number of independent loci available provides both power and accuracy and allows for the separation of real evolutionary effects from noise. This remains true even when considering just a few individuals, making genomic approaches much more versatile than more traditional methods utilising a few loci and large numbers of individuals (McMahon et al. 2014).

Population genomic analyses have a wide variety of applications and can be used to accurately gauge population structure and connectivity, taxonomic relationships, genetic diversity, and the demographic history of a species (Shafer et al. 2015; Allendorf et al. 2010; Steiner et al. 2013). Despite the recent rise in the number of genomic sequences obtained, many of these studies are still restricted to already well-studied species such as humans, model organisms and domesticates. However, the use of population genomic analyses is often extremely useful in solving questions about the evolutionary history of less studied, yet evolutionarily important lineages.

One little studied but ecologically important lineage is the family Hyaenidae (hyenas and aardwolf). Hyaenidae occupy a species-poor (four extant species) branch within the Feliformia suborder in a sister position to the Felidae family. However, despite its close relationship to the well-studied Felidae family, of which a number of genomes have already been sequenced (Abascal et al. 2016; Dobrynin et al. 2015; Cho et al. 2013), very few genetic studies have been carried out on the Hyaenidae family. Members of Hyaenidae occupy a variety of different niches. The most notable and arguably most important niche being that of the scavenger (Gusset and Burgener 2005; Watts and Holekamp 2007). Scavengers are known to be important for maintaining healthy ecosystem function with profound roles in nutrient cycling and influencing disease dynamics (Benbow et al. 2015).

The brown hyena (*Parahyaena brunnea*) is predominantly a scavenger, mainly feeding on large vertebrate carrion (Watts and Holekamp 2007). It is generally found in arid areas across southern Africa and is listed as ‘Near Threatened’ by the International Union for Conservation of Nature (IUCN). The brown hyena is the rarest of all extant hyena species with estimates of population size being less than 10,000 individuals worldwide (Wiesel 2015). Despite its listing as Near Threatened, brown hyenas continue to be persecuted, often considered as problem animals by farmers or killed for trophy hunting. Incidental and often deliberate poisoning, shooting and trapping of these animals all hamper the survival of this ecologically important species (Kent and Hill 2013).

Genetic studies of the brown hyena are very limited but have hinted towards very low genetic diversity within the species (Rohland et al. 2005; Knowles et al. 2009). Species-wide genetic comparisons using a short fragment of the mitochondrial cytochrome b gene found no variability in this region regardless of sample origin (Rohland et al. 2005). Moreover, a study investigating population structure within Namibian brown hyena, utilising microsatellites, also found no detectable population structure (Knowles et al. 2009). The inability of both of these studies to find population structure perhaps stems from very low levels of genetic diversity within the brown hyena. These early indications of low diversity are important to follow up on and investigate. Even though genetic diversity does not necessarily correlate with current day population sizes (Bazin et al. 2006; Leffler et al. 2012), it is still an important factor in understanding past evolutionary events. Importantly, knowledge of the evolutionary processes affecting a species is critical to inform conservation plans aimed at the long term management of its evolutionary potential (Romiguier et al. 2014).

Here we present complete mitochondrial and nuclear genomic analyses of brown hyena from populations across its range in Southern Africa (Fig. 1). We find both mitochondrial and nuclear genomic diversity to be extremely low, lower than in any other mammalian species published to date. Our data suggest that this low diversity results from a continuous decrease in effective population size over the last million years. Moreover, our low-coverage genomes reveal a number of potential sub-populations of brown hyena across its range.

**Figure 1:**
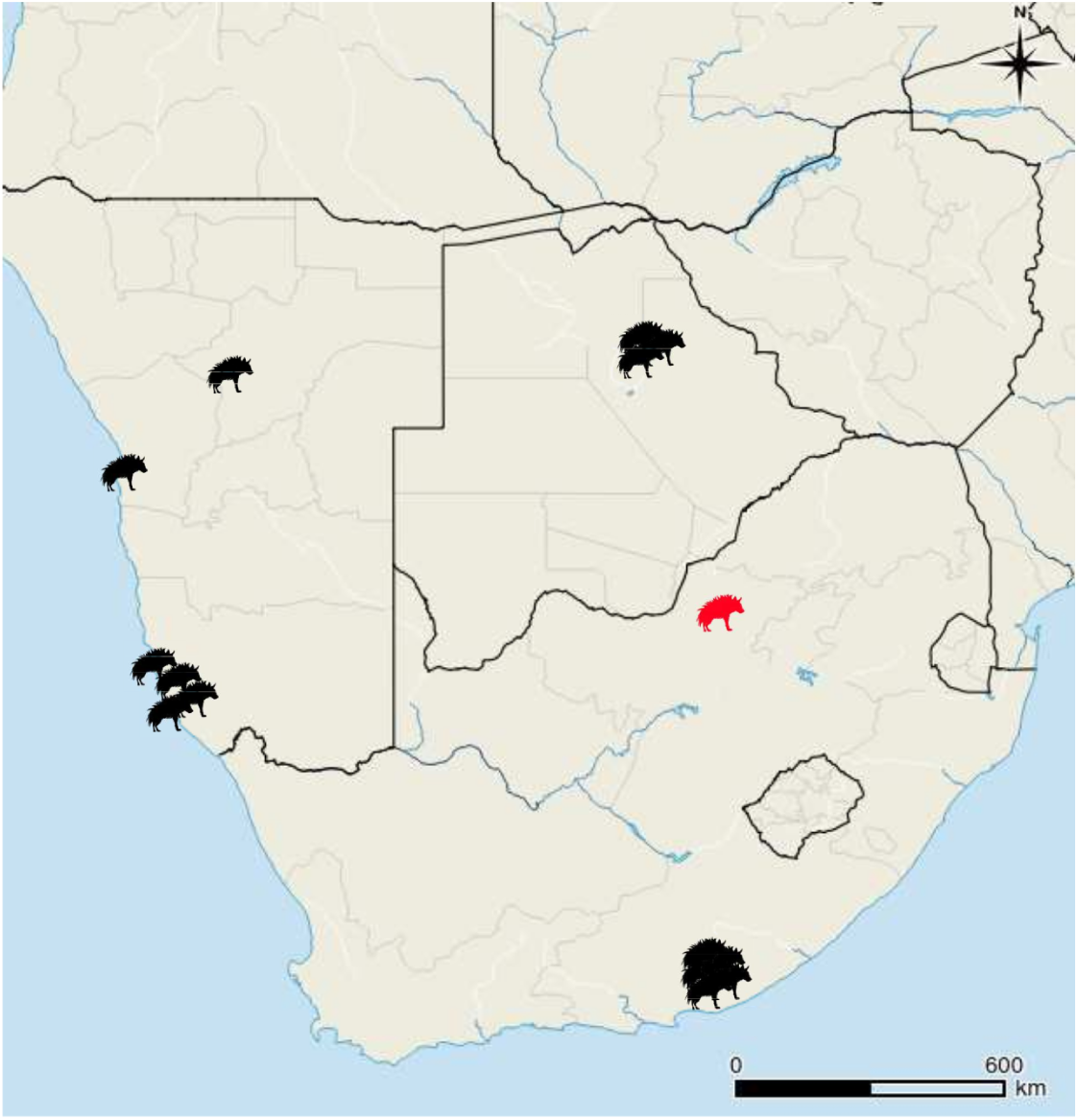
Map of the sampling locations of the wild-caught brown hyena included in this study. The red hyena indicates the original area of the South African population prior to translocations in 2003.

## Results

### Genome reconstructions

A *de novo* assembly of a striped hyena nuclear genome with a scaffold N50 of ~2Mbp was assembled using Allpaths LG (Butler et al. 2008), default parameters and an additional gap closing step using Gapcloser (Luo et al. 2012) (Supplemental Table S1). BUSCO analyses (Simão et al. 2015) using both the eukaryotic and mammalian databases show high levels of complete BUSCOs (Supplemental Table S2) giving us confidence that our assembly is of good quality and completeness. With this as reference, we successfully mapped low coverage nuclear genomes (2.1 - 3.7x) from 14 wild-caught brown hyena individuals originating from Namibia, South Africa and Botswana and a high coverage nuclear genome from a captive individual (Supplemental Tables S3 and S4).

Due to the lack of a published brown hyena mitochondrial genome, we assembled the complete *de novo* mitochondrial genome from a captive individual using MITObim (Hahn et al. 2013). We used default parameters apart from mismatch value, where we used zero, and three different bait reference sequences (domestic cat (U20753.1), spotted hyena (JF894377.1) and striped hyena (NC_020669.1)). All three independent MITObim runs produced identical brown hyena mitochondrial sequences, suggesting that our reconstructed mitochondrial genome is correct. We then mapped the remaining wild samples to this sequence (Supplemental Table S4).

### Genetic diversity

Mitochondrial DNA diversity estimates (Fig. 2) using the 14 wild caught mitogenomes from our data set showed that the brown hyena has the lowest diversity when compared to a number of other mammalian mitochondrial genomes as analysed in previous studies (Dobrynin et al. 2015; Miller et al. 2011). All brown hyena individuals shared an average of only three mutational mismatches (k) among them. This is about three times lower than the species with the next lowest level of mitochondrial diversity, the tasmanian devil (*Sarcophilus harrisii*), with a k value of 10, and 30 times lower than the gorilla (consisting of both *Gorilla gorilla* and *Gorilla beringei*), which has a k value of 92.

**Figure 2:**
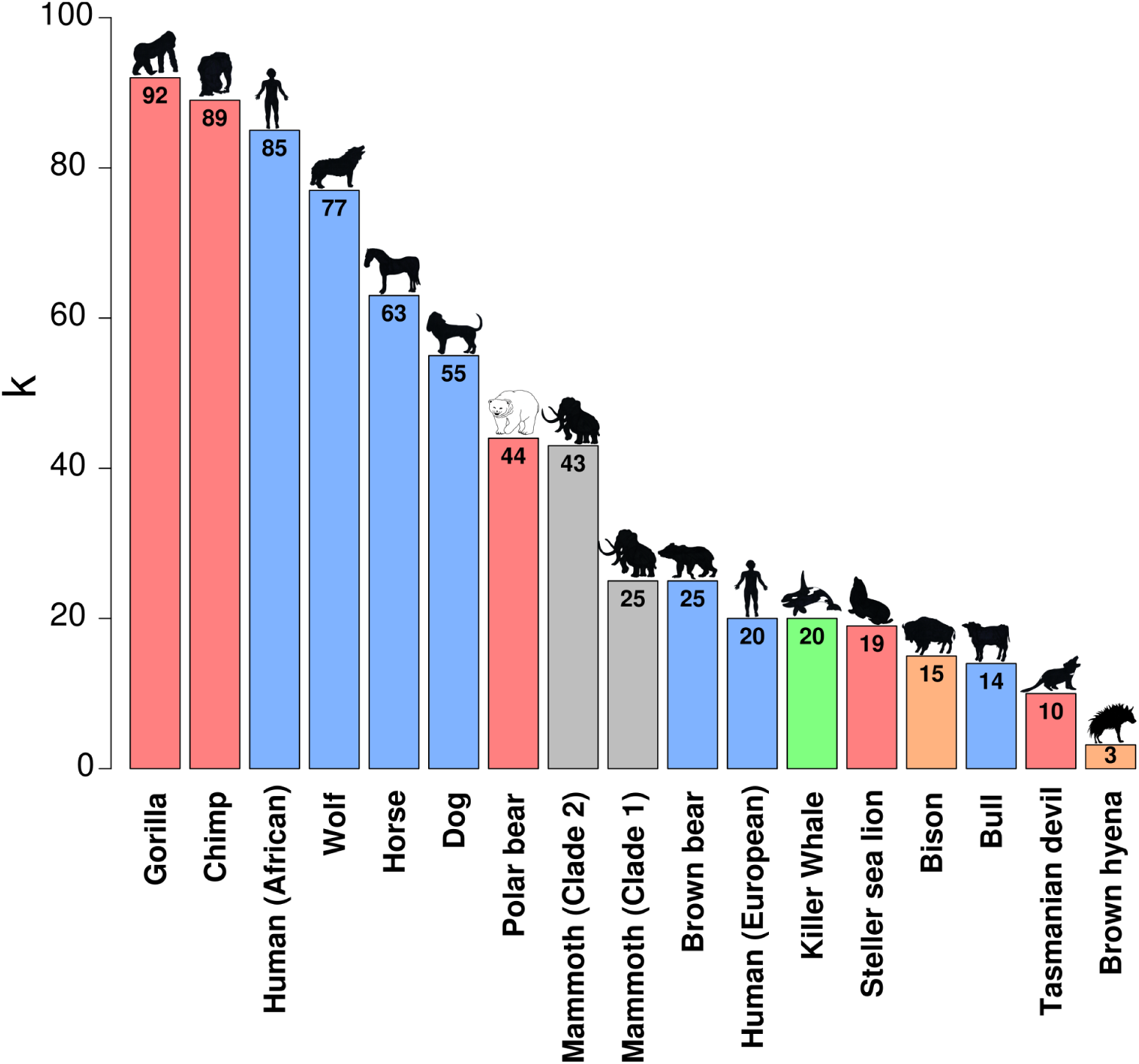
Mitochondrial diversity comparisons of the brown hyena to a number of other species/populations for which such data are available (Dobrynin et al. 2015; Miller et al. 2011). k represents the average number of substitutions expected between two randomly selected individuals of the same population, bar colours represent conservation status (red - endangered, grey - extinct, orange - near threatened, blue - least concern, green - unavailable) according to the IUCN.

We also calculated genome-wide nuclear heterozygosity estimates of our high coverage captive individual as these are considered a good proxy for nuclear genomic diversity. Results show the brown hyena to have the lowest of any species included in this study, most of which are considered “endangered” (Fig. 3A). These results are consistent with the very low levels of diversity we found within the mitochondrial genome.

**Figure 3:**
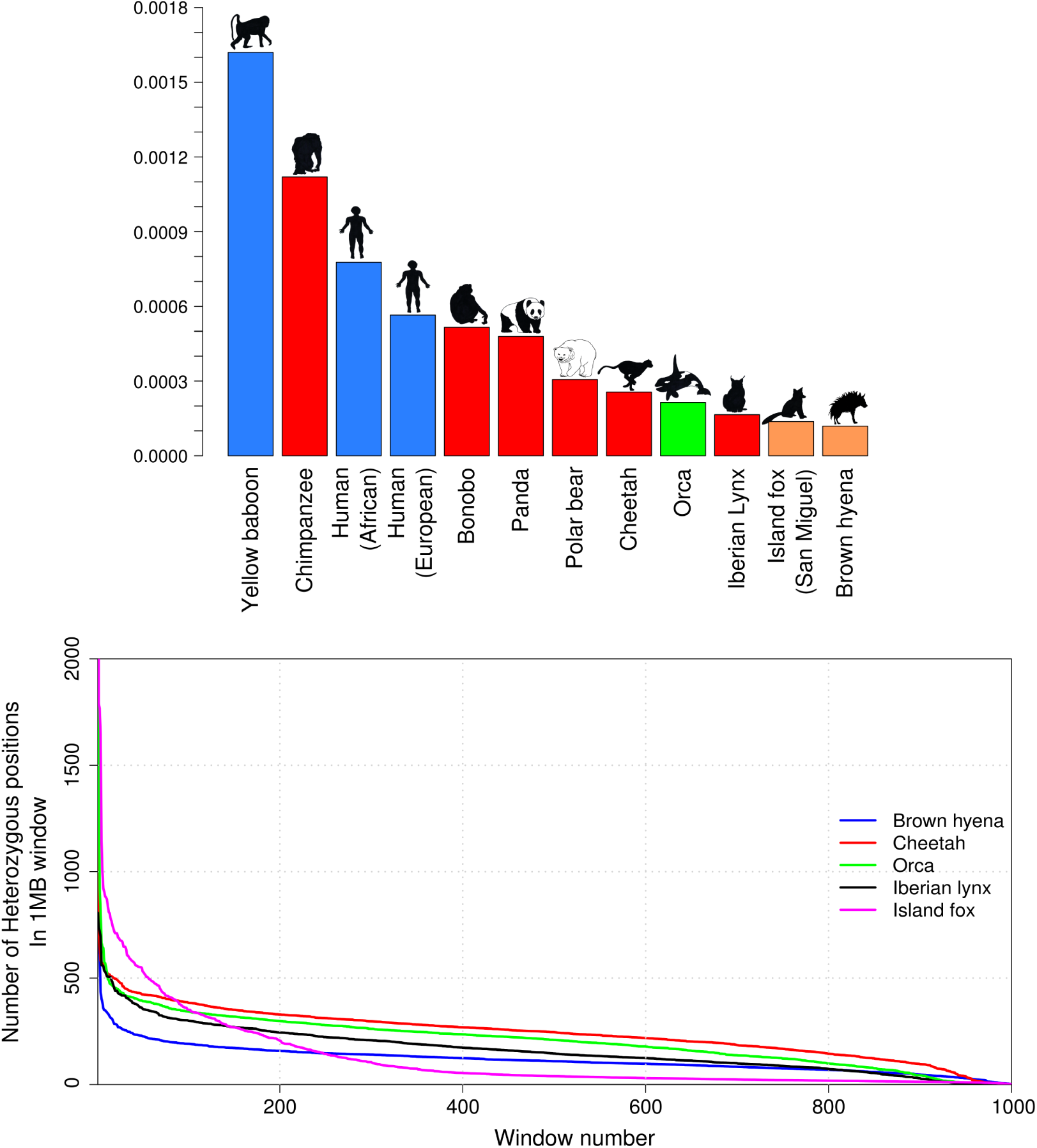
Estimated heterozygosity levels in the brown hyena and a comparison to other mammalian species. **A)** Average genome-wide heterozygosity comparisons. Y axis represents the average proportion of sites within the nuclear genome to be heterozygous. Bar colours represent conservation status (red - endangered, orange - near threatened, blue - least concern, green - insufficient data) according to the IUCN. **B)** Heterozygosity density comparisons between the four species with the lowest estimated average genome-wide heterozygosity levels in this study. Y axis represents the number of heterozygous sites within the 1Mbp window. X axis represents the window. Colours represent species ( blue - brown hyena, black - Iberian lynx, green - orca, red - cheetah, magenta - Island fox).

The brown hyena individual sequenced to high coverage is known to have been born from wild-caught parents, albeit in a captive environment, and should not display large amounts of inbreeding that can be found in captive bred populations. However, to investigate signs of inbreeding, we measured levels of heterozygosity in various window sizes across the brown hyena nuclear genome. The results show no considerable stretches of homozygosity (Supplemental Fig. S1), suggesting that there are no significant signs of inbreeding. We further analysed the distribution of heterozygosity across the genome by randomly selecting 1000 non-overlapping windows of 1Mbp in size and comparing the results to the four other species with the lowest heterozygosity included in this study, the orca (*Orcinus orca*), cheetah (*Acinonyx jubatus*), Channel Island fox (*Urocyon littoralis*) and Iberian lynx (*Lynx pardinus*) (Fig. 3B). The brown hyena genome has consistently lower levels of heterozygosity across the genome when compared to the Iberian lynx, orca and cheetah, excluding the windows with the lowest levels of heterozygosity which may indicate higher levels of inbreeding in the other species. However, it can be seen that, while the brown hyena has a lower level of average genome-wide diversity, for the majority of the windows, the Channel Island fox has lower heterozygosity than the brown hyena. Regions of high levels of heterozygosity amongst a sea of low heterozygosity have been previously reported for the Channel Island fox (Robinson et al. 2016). Moreover, it should also be noted that as each estimate only shows the genomic diversity of a single individual and levels are expected to vary within natural populations, some caution must be taken when interpreting these results.

### Demographic history

As sex chromosomes are known to have a different demographic history than autosomes, we found and removed scaffolds in our assembly related to the X chromosome before running demographic analyses. We aligned our striped hyena assembly by synteny using Satsuma (Grabherr et al. 2010) to the cat X chromosome (126,427,096bp) and found 195 scaffolds (Supplemental Table S5), totaling 117,479,157bp. The alignment was visualised using Circos (Krzywinski et al. 2009) (Supplemental Fig. S2). The Circos alignment shows that the scaffolds found using Satsuma cover the complete cat X chromosome. These scaffolds were then removed before running a Pairwise Sequentially Markovian Coalescent (PSMC) model (Li and Durbin 2011). PSMC analyses using the brown hyena autosomes were consistent with its low genomic diversity (Fig. 4; Supplemental Fig. S3) and revealed a continuous gradual decrease in effective population size over the last one million years. This was then followed by a more rapid recent decrease in effective population size at the end of the Late Pleistocene.

**Figure 4:**
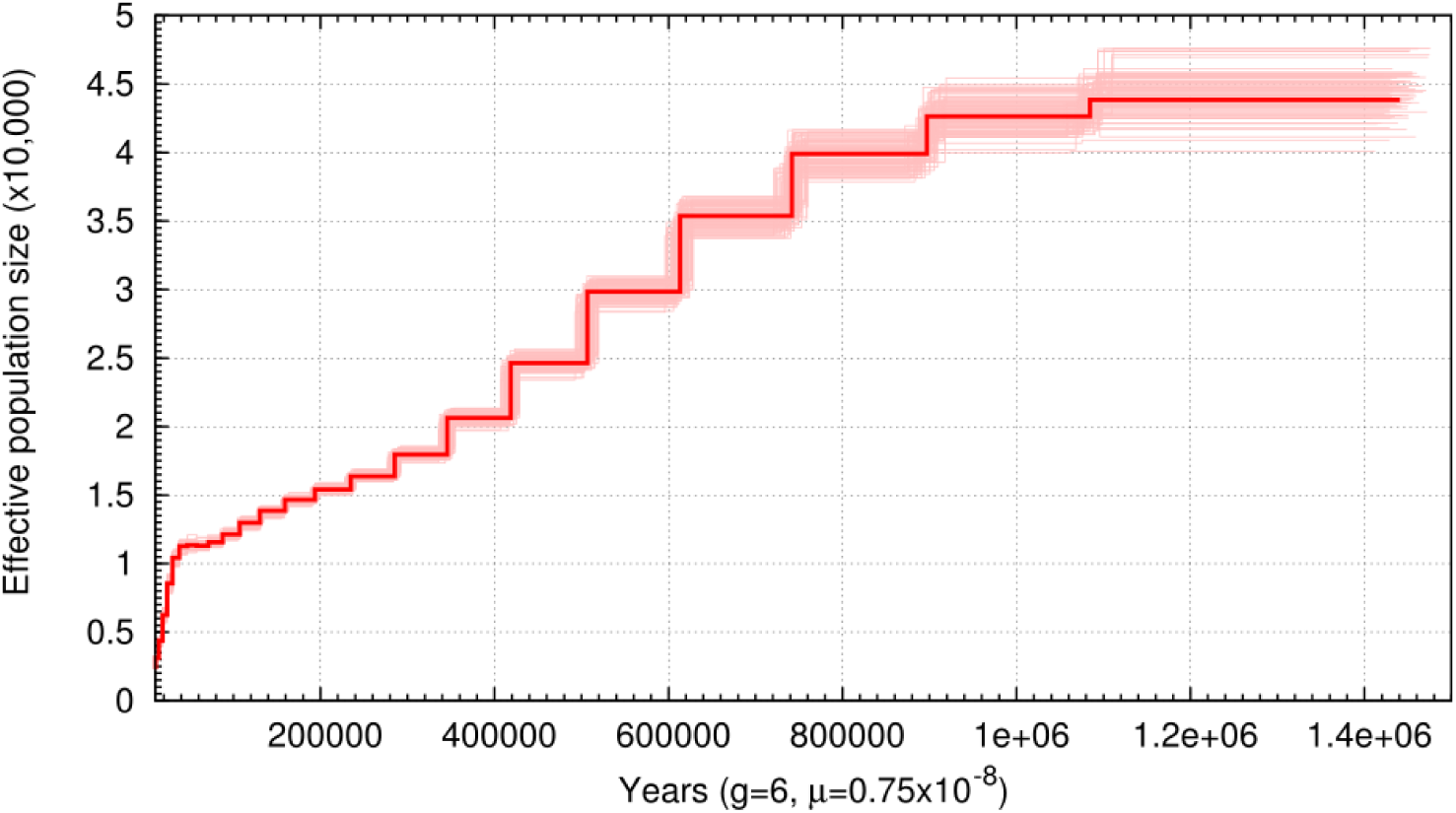
Pairwise Sequentially Markovian Coalescent (PSMC) model plot of the autosomes of one high coverage brown hyena. The Y axis represents effective population size and the X axis represents time in years. Light red bars show bootstrap support values. Calibrated using a generation time (g) of 6 years and a mutation rate (μ) of 7.5×10-9 per generation.

### Population structure

In order to infer potential population structure, we reconstructed a haplotype network using the mitochondrial genomes of the wild-caught individuals. This network revealed some phylogeographic structure with all but one haplotype being geographically restricted. This one shared haplotype was shared among individuals from all three sampled countries (Fig. 5A).

**Figure 5:**
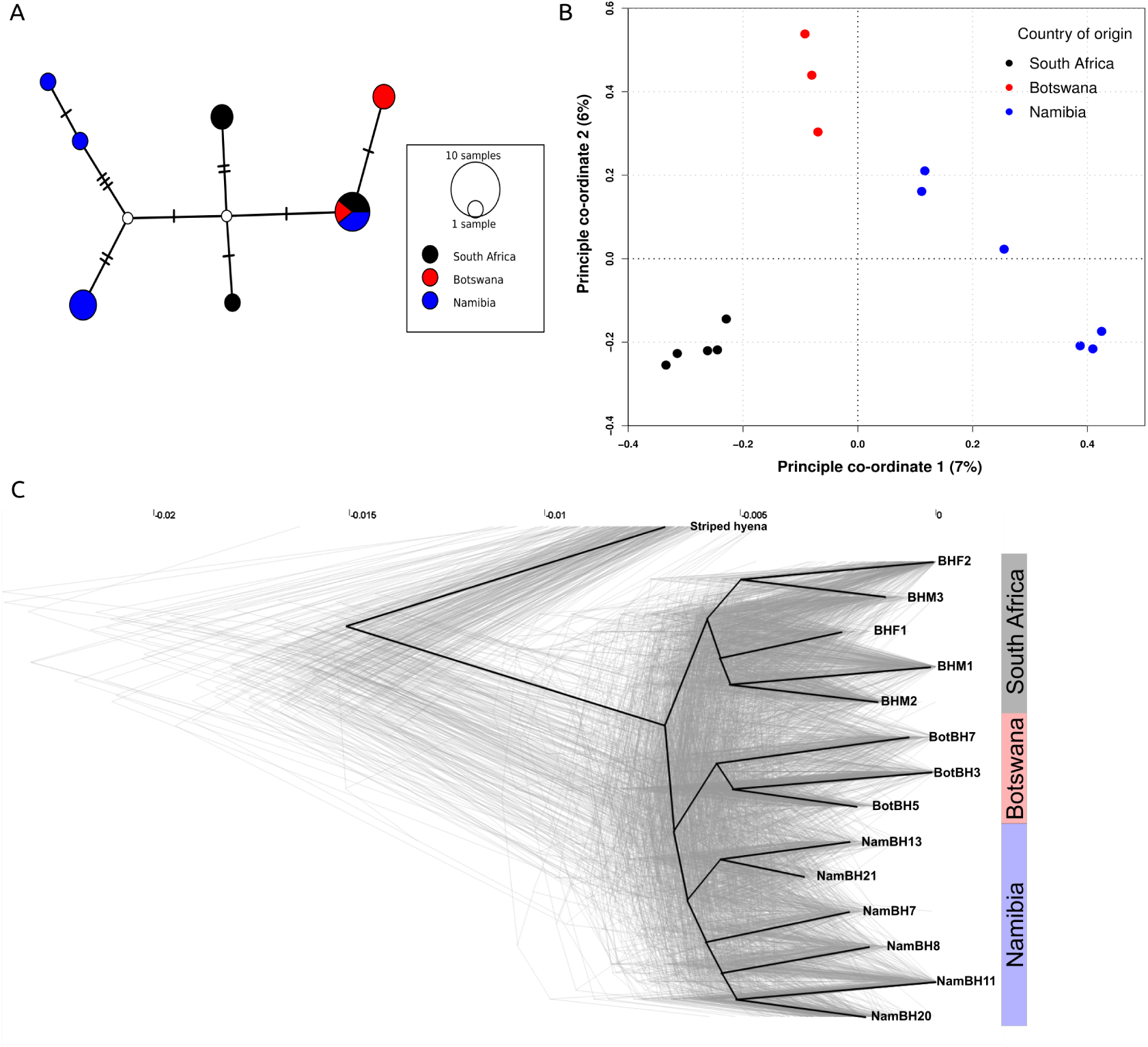
Genomic diversity analyses **A)** Median joining haplotype network of the 14 wild-caught brown hyena individuals included in this study. Lines on the connecting branches represent single base differences, size of the circle represents number of individuals belonging to a single haplotype and colours represent sampling country (black - South Africa, blue - Namibia, red - Botswana). **B)** Principal components analysis produced using genotype likelihoods for the low coverage genomes of the 14 wild-caught brown hyena individuals in this study. Colours represent country of origin (black - South Africa, blue - Namibia, red - Botswana). Percentages on the X and Y axis represent the percentage of variance explained by each respective component. **C)** Densitree phylogenetic tree. Light grey lines represent phylogenetic trees produced from single scaffolds. The dark black line represents the root canal as defined by Densitree.

We then used the mapped genomes of the wild-caught individuals to infer nuclear population structure by carrying out principal component analyses (PCA) and maximum likelihood (ML) phylogenetic analyses. The results showed that clusterings or clades of individuals were in all cases consistent with the geographical origin of the individuals (Fig. 5B; Supplemental Fig. S4). PCA analyses using both single base identity by state (IBS) and genotype likelihoods (GL) produced similar results (Fig. 5B; Supplemental Fig. S5).

In contrast to whole nuclear genomic data, both PCA and phylogenetic analyses using single scaffolds did not produce unanimous phylogeographic results. Independent PCA results for the nine longest scaffolds can be seen in Supplemental Fig. S6. Although they do generally support some phylogeographic structure, individuals from different regions partially intermingle in these plots. When running independent per-scaffold ML phylogenetic analyses with scaffolds over 2MB, we find that in 49 out of 333 trees, the South African samples form a monophyletic clade, in 72 out of 333 trees the Botswana samples form a monophyletic clade and in 17 out of 333 trees the six Namibian samples form a monophyletic clade. This non-unanimous pattern shows up as clouds surrounding all nodes within the brown hyena lineage when visualising all independent ML trees simultaneously using Densitree (Bouckaert 2010) (Fig. 5C). Although few individual trees support monophyly of either of the three geographical groups, the root canal (consensus tree with highest clade support), defined by Densitree, nevertheless shows a similar topology as the tree constructed using the complete nuclear genome (Fig. 5C; Supplemental Fig. S4). Importantly, it is again fully consistent with the geographical origin of the individuals.

Admixture and population structure analyses using NGSadmix (Skotte et al. 2013) reached convergence for K values of two to five, indicating that there may be anywhere from two to five different populations inhabiting southern Africa (Fig. 6). When considering a K value most congruent with the number of countries of origin, K3 (Fig. 6B), individuals NamBH13 and NamBH21 stand out as being “admixed” between all three populations. This could be a byproduct of the analysis trying to find a population for them. A K value of four however, shows them as an independent population (Fig. 6C). The latter is the most parsimonious result as it is consistent with their geographic origins (northern Namibia while all other Namibian samples are from the south) and previous findings investigating the distribution patterns of individuals across Namibia (Supplemental Fig. S7).

**Figure 6:**
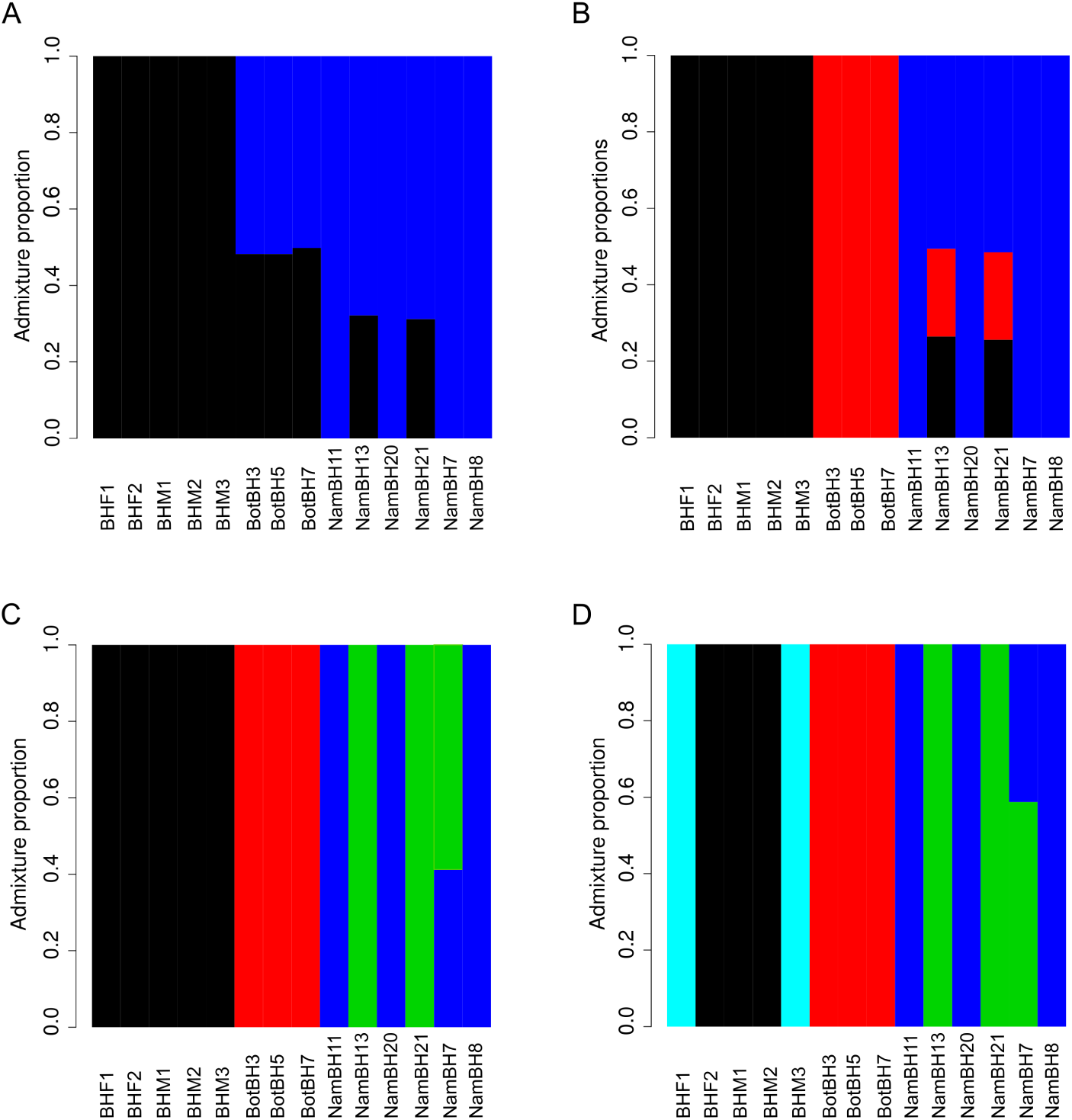
Admixture plots produced using different K values in NGSAdmix. **A)** K2 **B)** K3 **C)** K4 **D)** K5. Yaxis represents the admixture proportion and x axis represents the individual.

We carried out D-statistics comparisons testing for population structure (Fig. 7). A high D-value could either represent differential levels of admixture or more recent common ancestry and therefore an incorrect predefined topology. Taking the latter into account, we placed individuals into one of three (defined as: Namibia, Botswana, or South Africa) or one of four (defined as: northern Namibia, southern Namibia, Botswana, or South Africa) predefined populations and compared the D values produced from “correct” topologies, i.e branches H1 and H2 belong within the same population (e.g. Fig. 7A), and “incorrect” topologies, i.e branches H2 and H3 belong within the same population (e.g. Fig. 7B). When considering four populations (Fig. 7C) it can consistently be seen that when we break the predefined population structure and therefore topology, we find a higher D value than when individuals in the same predefined population are in the H1 and H2 positions. Assuming that the correct topology is that with the lowest D value and as there is clear separation between D values recovered when breaking and not breaking the predetermined population structure when using four predefined populations, we conclude that there are indeed four populations within our dataset. This pattern is not seen when only three predefined populations (Fig. 7D) are considered as there are many overlapping D values within the Namibian population when “correct” and “incorrect” topologies are tested. This led us to reject a possibility of one single population within Namibia. These results suggest four populations, with a split between northern and southern Namibia. This analysis revealed the same population structure as the NGSadmix analyses, thus corroborating the observation that this dataset consists of four populations rather than three.

**Figure 7:**
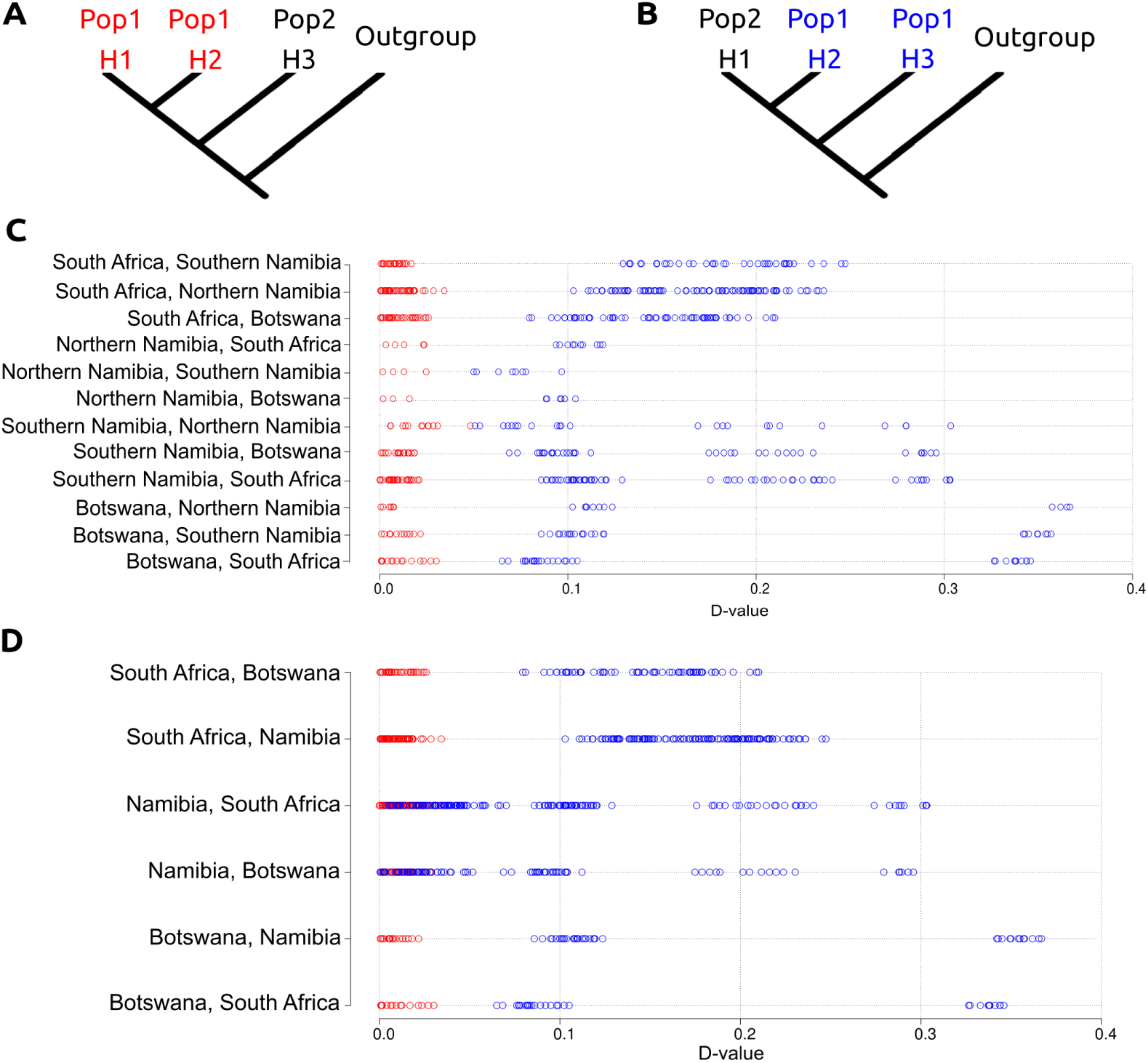
Topology test using D-statistics. **A)** D-statistic analysis demonstrating the correct predetermined population structure. **B)** D-statistic analysis demonstrating the incorrect predetermined population structure. **C)** D-statistics comparisons when four “populations” are determined a priori. **D)** D-statistics comparisons when three “populations” are determined a priori. Red coloured circles represent comparisons when the predefined population structure is not broken within the comparison. Blue coloured circles represent comparisons when the predefined population structure is broken within the comparison. X axis represents the D-value.

We carried out IBS analyses on a number of other mammalian species for which species-wide low coverage data was available in order to compare these with the population structure within the brown hyena. Each species, including the brown hyena, had 10 representatives randomly selected and pairwise distance comparisons were carried out within each species (Supplemental Figure. S8). For the brown hyena, most pairwise distance values clustered together, representing a relatively high level of shared diversity between different putative populations and suggesting extensive gene flow among these populations. This pattern was not as strong, however, as in some species, such as domesticated sheep which are known to be fairly panmictic (Peter et al. 2007) and still suggests some level of differentiation within the brown hyena. We also found no abnormalities, i.e. potential subspecies, large differences between populations or indications that a number of the individuals were extraordinarily related which could be driving the “population” signal we find in the brown hyena.

## Discussion

Hyaenidae occupy a major, albeit species poor branch within Feliformia. The family has a rich fossil history but is now restricted to only four extant species (Koepfli et al. 2006). The abundance of fossil specimens has led to a number of studies focused on the taxonomic relationships within fossil Hyaenidae (Werdelin and Solounias 1991; Turner et al. 2008). Little attention, however, has been focused towards the evolution and population status of the extant species, especially on a molecular level. Despite the brown hyenas rarity, molecular analyses of the brown hyena are currently limited to two studies on microsatellites and short regions of mitochondrial DNA (Rohland et al. 2005; Knowles et al. 2009). Both hinted at low genetic diversity but had very limited results with regard to population structure.

### Genomic diversity

Using complete nuclear and mitochondrial genomes of 14 wild-caught brown hyena originating from across southern Africa, brown hyenas displayed the lowest level of both mitochondrial and nuclear genomic diversity when compared against other mammalian species for which comparable data were available, including many endangered species (Fig. 2; Fig. 3A). Genomic diversity was even lower than in species famously known to have extremely low levels of genomic diversity, such as the cheetah (Dobrynin et al. 2015) and the Iberian lynx (Abascal et al. 2016), both of which have gained considerable research attention because of this characteristic. In addition to finding very low levels of genomic diversity, demographic analyses of the brown hyena show a gradual, yet steady decline in effective population size over the last million years, with a more rapid decline towards the end of the Late Pleistocene (Fig. 4). The brown hyena is known to have once had a more extensive range, with Middle Pleistocene fossils having been found in Kenya (Werdelin and Barthelme 1997). The continuous decrease in effective population size and low levels of genomic diversity seen today may have occurred with the shrinking of suitable habitats during the Pleistocene (deMenocal 2004) and potentially coincided with the migration of new competitors, such as jackals, into Africa (Koepfli et al. 2015).

Interestingly, despite very low levels of genetic diversity within both the nuclear and mitochondrial genomes, we found no strong signs of inbreeding in the nuclear genome (Fig. 3B; Supplemental Fig. S1). This was an unexpected result as low genomic diversity is generally expected to arise from high levels of inbreeding (Willoughby et al. 2015). Low genomic diversity has commonly been associated with decreased fitness and higher extinction rates (Spielman et al. 2004; Reed and Frankham 2003). However, the lack of detectable inbreeding in our high coverage individual could mean that this species has evolved into a state of genetic stasis, allowing low genetic diversity to persist across the genome, not strongly influencing the survivability of the species. This genetic stasis could have evolved as a result of the slow but continuous decrease in effective population size seen over the last million years and lack of detectable bottlenecks during that time (Fig. 4). This slow continuous decrease could hamper the influence of genetic drift therefore allowing purifying selection to maintain variability in a number of loci potentially important for the survival of the species. The retention of relatively high levels of potentially important adaptive variation allowing for the retention of evolutionary potential has recently been found in Channel Island foxes (Robinson et al. 2016). Heterozygosity hotspots were shown to be present in a number of genes with high levels of ancestral variation, despite low levels of genome-wide heterozygosity in the Channel Island fox. However, as we see heterozygosity to be fairly evenly distributed across the genome, this doesn’t seem to be the case for the brown hyena and it has perhaps been able to persist despite the lack of adaptive variation due to some other factor. Further research would however be required to definitively know whether the brown hyena goes against the commonly perceived notion that high levels of adaptive variation are required to ensure a species survival.

### Brown hyena population structure

By using a combination of different population structure analyses on the entire nuclear genome we were also able to, for the first time, define population structure within the brown hyena (Fig. 5; Fig. 6 and Fig. 7) despite its very low levels of genetic diversity. Results were in concordance with geographic structuring. Furthermore, by comparing these results to those produced using mitochondrial genomes and single scaffolds, we show that even when using millions of loci (i.e. single scaffolds >2Mbp (Fig. 5C; Supplemental Fig S6)) difficulties with accurately defining population structure can arise. This maybe due to the very low genomic diversity within the brown hyena. This finding reinforces the value of using whole nuclear genomes in population analyses over other approaches such as microsatellites and reduced genomic representation techniques, especially in species suspected to have low genomic diversity.

### Conservation implications

Our findings provide a greater understanding into the evolution and population structure of the brown hyena, which has wider implications in aiding conservation of this species. Due to the increase in agriculture, many habitable areas for the brown hyena have now been fragmented, and the connectivity between populations has therefore been reduced. The wide geographical range of the brown hyena, coupled with the extremely low levels of genomic diversity, indicates that the brown hyena persists at low population densities, which is in concordance with previous studies analysing natural population densities (Kent and Hill 2013). This demonstrates the importance of connectivity within the species. Given the overall evidence for substantial gene flow between the different populations, well implemented translocations may certainly become important for the survivability of this species in the future. With better definitions of population boundaries, translocations can be implemented in such a way as to avoid outbreeding depression and increase the chances of an animal successfully adapting to its new environment (Allendorf et al. 2010). Moreover, previous studies have shown that translocated brown hyena are able to settle into their new environments and do not return to their original locations (Weise et al. 2015), and that human mediated translocations can lead to the rise of highly dense and successful populations (Welch and Parker 2016). These studies further demonstrate the ability of this species to adapt and accept translocations, and the utility these techniques could have in the future.

## Methods

### Samples

In total, 15 brown hyena samples were used for this study; five from South Africa, six from Namibia, three from Botswana (Fig. 1) and one from Tierpark Berlin (Supplemental Table S3). One female striped hyena (*Hyaena hyaena*) from Tierpark Berlin was also included to be used for reference based mapping.

### Striped hyena *de novo* assembly

We extracted DNA from Hyena2069, the Tierpark Berlin striped hyena sample, on a KingFisher Duo robot using the blood DNA extraction kit according to the manufacturer’s instructions. The extract was then built into two PCR free Truseq Illumina sequencing libraries, one with 180bp and one with 670bp short inserts. Two Nextera mate-pair libraries were also constructed with sizes of 3kbp and 6kbp. All of the above-mentioned libraries were constructed at the National Genomics Infrastructure (NGI) in Stockholm. All libraries were then sequenced on an Illumina HighseqX using 2x150bp paired-end sequencing at the NGI in Stockholm. The 180bp and 670bp insert libraries were sequenced on one lane each, whereas the mate-pair libraries were multiplexed and sequenced together on a single lane.

The NGI trimmed Illumina adapter sequences from the raw illumina reads using Trimmomatic (Bolger et al. 2014) and performed a *de novo* assembly using Allpaths LG (Butler et al. 2008) with default parameters. We then performed an additional gap closing step using Gapcloser (Luo et al. 2012). Assembly quality and completeness was assessed using BUSCOv2 (Simão et al. 2015) using both the eukaryote and mammalian BUSCO databases (Supplemental Table S2).

### Captive brown hyena sample

We extracted DNA from a single brown hyena sample from Tierpark Berlin on a KingFisher Duo robot using the cell and tissue DNA extraction kit. The extract was then built into a PCR free Truseq Illumina sequencing library using a 350bp insert size by the NGI in Stockholm. This library was then sequenced on an Illumina Highseq X using 2×150bp paired-end sequencing at the NGI in Stockholm.

### Wild-caught brown hyena samples

We extracted DNA from the six blood and three tissue samples using a DNeasy blood and tissue extraction kit, following the manufacturer’s protocol. We extracted the five hair samples using the DY04 user modified version of the DNeasy kit and protocol. We fragmented DNA extracts using a Covaris sonicator into ~500bp fragments. Fragmented extracts were then constructed into Illumina sequencing libraries using a modified version of the protocol set out by Meyer and Kircher (Meyer and Kircher 2010; Fortes and Paijmans 2015). Library molecules from 400bp to 900bp were selected using a Pippin Prep Instrument (Sage Science) and sequenced on an Illumina Nextseq 500 at Potsdam University, Germany.

### Raw data treatment

We trimmed Illumina adapter sequences and removed reads shorter than 30bp from the raw reads of the 15 brown hyena samples using Cutadapt v1.8.1 (Martin 2011) and merged overlapping reads using FLASH v1.2.1 (Magoč and Salzberg 2011).

### Mitochondrial genome reconstruction

As no brown hyena mitochondrial sequence was available, we reconstructed one using the shotgun data from our single high coverage individual. We assembled the mitochondrial genome through iterative mapping using MITObimv1.8 (Hahn et al. 2013) on 40 million trimmed and merged reads, subsampled using seqtk (Li 2012). We removed duplicate reads using prinseq (Schmieder and Edwards 2011). MITObim was performed in three independent runs using three different starting bait reference sequences. The references included the domestic cat (U20753.1), spotted hyena (JF894377.1) and striped hyena (NC_020669.1). We implemented MITObim using default parameters apart from mismatch value where we used zero. Output maf files were converted to sam files and visualized using Geneious v9.0.5 (Kearse et al. 2012). Consensus sequences were constructed in Geneious using a 75% base call consensus threshold, and only sites with over 20x coverage were considered.

The reconstructed mitochondrion served as a reference sequence for subsequent mitochondrial mapping analyses. We mapped the trimmed and merged reads from our 14 wild brown hyenas to the reconstructed reference sequence using BWAv0.7.15 (Li and Durbin 2009), using the mem algorithm and default parameters and parsed the mapped files using Samtools v1.3.1 (Li et al. 2009). The consensus sequences were constructed using ANGSDv0.913 (Korneliussen et al. 2014), only considering reads and bases of quality scores greater than 25.

### Mitochondrial analyses

The mitochondrial genomes from the 14 wild-caught brown hyena were aligned using Mafftv7.271 (Katoh and Standley 2013). We constructed a median joining haplotype network of the alignment using Popart (Leigh and Bryant 2015). Nucleotide diversity (pi = 0.000140667) was estimated using Popart. The k value was calculated by multiplying the pi value by the total length of the brown hyena mitochondria. This number was then compared against those from previous studies (Dobrynin et al. 2015; Miller et al. 2011) (Fig. 1B).

### Low coverage nuclear genome analyses

Trimmed and merged data were mapped to the *de novo* striped hyena assembly using BWA v0.7.15 (Li and Durbin 2009) and parsed using Samtools v1.3.1 (Li and Durbin 2009). We applied the following filtering options for all analyses involving ANGSD (Korneliussen et al. 2014): we only considered sites where at least 10 individuals had coverage (-minInd 10), only included sites for which the per-site coverage across all individuals was less than 75. We implemented quality filtering by setting a minimum base quality score of 25 (-minQ 25), minimum mapping quality score of 25 (-minMapQ 25) and only allowed reads that mapped uniquely to one location (-unique_only 1). We also adjusted quality scores around indels (-baq 1) (Li 2011).

### Brown hyena population structure

Principle component analyses (PCA) were carried out using both single read IBS analyses and GL analyses in ANGSDv0.913 (Korneliussen et al. 2014). IBS analyses were restricted to SNPs occurring in at least two individuals. This was done to remove singletons which could represent sequencing errors. We computed genotype likelihood in ANGSD and converted outputs to a covariance matrix using ngsTools (Fumagalli et al. 2014). Covariance matrices were converted into PCA outputs and visualized using R (R Development Core Team 2008). For the phylogenetic analyses, we performed Maximum likelihood analyses with Raxml v8.2.10 (Stamatakis 2014), specifying the striped hyena as outgroup and using the GTR+GAMMA substitution model. We prepared the infile for this by computing consensus sequences using ANSGD with the above-mentioned filters. We then performed genome-wide alignments, removed sites with missing data in three or more individuals, sites where singletons occurred within the brown hyena ingroup and invariant site positions using a custom Perl script.

We then repeated the phylogenetic and IBS PCA analyses using single scaffolds. PCA analyses were carried out using nine independent analyses (scaffolds 0-8) and Maximum likelihood analyses were carried out independently for single scaffolds with a length larger than 2MB. We calculated admixture proportions using NGSadmix (Skotte et al. 2013) setting K values from 2-7. We used ANGSD genotype likelihood values as input, only including SNPs with a p value of less than 1e-6. NGSadmix analyses were repeated a maximum of 100 times per K. Only those that converged (produced a consistently identical likelihood score) within these 100 analyses were considered as meaningful. D-statistic analyses were implemented in ANGSDv0.913, sampling a single base per site while specifying the striped hyena as outgroup with default parameters.

### Comparative population structures

In order to compare the population structure within the brown hyena to those of other species, 10 individuals per species were randomly selected for a number of different mammalian species for which such data was publically available (Supplemental Table S6). Comparisons between individuals were performed using single base IBS, only considering sites where at least 7 individuals had coverage and SNPs that occurred in at least 2 individuals. Other filtering options included; a minimum base quality score of 25 (-minQ 25), minimum mapping quality score of 25 (-minMapQ 25) and reads that mapped uniquely to one location (-unique_only 1). Quality scores around indels were also adjusted for (-baq 1).

### Species heterozygosity estimates

High coverage, single individual representatives for a number of species were assessed for heterozygosity levels. Raw data were selected from a range of different species (Supplemental Table S7). Raw reads were all treated comparably, using Cutadapt v1.8.1 (Martin 2011) to trim Illumina adapter sequences and FLASH v1.2.1 (Magoč and Salzberg 2011) to merge overlapping reads. We mapped each species to its respective reference sequence using BWAv0.7.15 (Li and Durbin 2009) and processed the mapped reads further using Samtools v1.3.1 (Li et al. 2009). To adjust for biases introduced by unequal levels of coverage, the resulting bam files were all subsampled to 20x using Samtools (Li et al. 2009). The folded site frequency spectrum and therefore heterozygosity was then calculated from sample allele frequencies, taking genotype likelihoods into account for each species representative calculated using ANGSDv0.913 (Korneliussen et al. 2014). For the 1Mbp window analysis, we used the same analysis to estimate heterozygosity as before but performed the analysis in non-overlapping windows of 1MB. We also removed any scaffolds less than 1Mbp in length.

### Demographic inference

The demographic history of the brown hyena was calculated using PSMC (Li and Durbin 2011), considering only the autosomal chromosomes. Scaffolds representing the X chromosome of the striped hyena were determined through a synteny analysis to the cat X chromosome (CM001396.2) using Satsuma synteny (Grabherr et al. 2010). These scaffolds were then removed along with any scaffold shorter than 1Mbp. A consensus diploid sequence was constructed using Samtools (Li et al. 2009) to be used as input for PSMC. PSMC was implemented using parameters previously shown to be meaningful when considering human data. 100 bootstrap analyses were undertaken. When plotting, we used a generation time of 6 years and a mutation rate of 7.5×10-9 per generation for autosomes.

In order to estimate the mutation rate, we carried out a pairwise distance analysis on the striped and brown hyena’s autosomes using a consensus base IBS approach in ANGSDv0.913. We then calculated the average per generation mutation rate assuming a divergence date of the two species to be 4.2mya (Koepfli et al. 2006), a genome-wide strict molecular clock and a generation time of 6 years. Additional analyses utilising different mutation rates based on the 95% confidence interval of the brown and striped hyena divergence (2.6mya and 6.4mya) can be seen in Supplemental Fig. S3.

### Data access

Raw sequencing reads can be found under the accession codes XXXXXX. The striped hyena nuclear genome assembly can be found at XXXXXX and the brown hyena mitochondrial genomes can be found at XXXXXX.

## Acknowledgements and funding

This work was supported by the European Research Council (consolidator grant GeneFlow # 310763 to M.H.). The authors also acknowledge support from Science for Life Laboratory, the Knut and Alice Wallenberg Foundation, the National Genomics Infrastructure funded by the Swedish Research Council, and Uppsala Multidisciplinary Center for Advanced Computational Science for assistance with massively parallel sequencing, as well as *de novo* assembly of the striped hyena and access to the UPPMAX computational infrastructure. L.D. acknowledges support from the Swedish Research Council and FORMAS. We would like to thank Prof. Yoshan Moodley for his suggestions on the manuscript. We would finally like to thank Binia De Cahsan for producing the animal icons found in Figures 1, 2 and 3.

## Author Contributions

The project was conceived by M.W. and M.H. M.W. and L.D. performed lab work; M.W., S.H., and A.B. performed DNA analyses and interpretation of results. I.W., V.L., R.W, D.M.P., F.S., and A.L. assisted with locating and sampling of specimens. Final editing and manuscript preparation was coordinated by M.W. All contributing authors read and agreed to the final manuscript.

